# Quasineutral dynamics shape coexistence and clearance in a model of *in vitro* phage–bacteria interactions

**DOI:** 10.64898/2025.12.25.696485

**Authors:** Jesús Ait Idir, Fatima Drubi, Marcos Rodríguez, Irene Cantallops, Pilar Domingo-Calap, Josep Sardanyés

## Abstract

Phage therapy, which uses viruses that infect bacteria to target and lyse specific bacterial pathogens, has re-emerged as a promising strategy to combat antibiotic-multiresistant bacteria. Advances in metagenomics and synthetic biology, together with systems biology approaches combining mathematical modeling with experimental data, provide excellent opportunities to understand phage-bacteria dynamics. Here we analyze a mathematical model successfully calibrated using *in vitro* data on the multidrug-resistant bacterium *Klebsiella pneumoniae* in the presence of the phage vB Kpn 2-P4. The model describes a system with a susceptible bacterial population that can generate phage-resistant mutants. By analyzing the equilibria and bifurcations of the model, we identify a coexistence scenario between phage-resistant bacteria and phages governed by a quasineutral line of equilibria with both stable and unstable segments. Biologically, this quasineutral structure implies that phage-resistant bacteria can persist across a wide range of phage densities without selective pressure favoring a unique outcome, making clearance highly sensitive to additional mortality mechanisms. The clearance of phage-resistant bacteria can be achieved by combining phage activity with an increased death rate of the resistant strains. This process is governed by a global transcritical bifurcation of the quasineutral line. Our model offers mechanistic insight into potential scenarios leading to the complete elimination of phage-resistant bacteria.

**Author summary:** Bacteriophages—viruses that infect bacteria—are being reconsidered as alternatives to antibiotics, but bacterial resistance to phages often emerges rapidly. Understanding when phages and bacteria coexist and when resistant bacteria can be eliminated remains a major challenge. In this study, we analyze a mathematical model calibrated with *in vitro* data describing interactions between bacteria, bacteriophages, and phage-resistant mutants. Using tools from dynamical systems theory, we show that coexistence between phages and resistant bacteria is organized by a quasineutral line of equilibrium states rather than a single stable outcome. This structure explains why long-term dynamics can be highly sensitive to initial conditions without being chaotic. We further identify a bifurcation that leads to the extinction of resistant bacteria when their effective mortality exceeds a critical threshold. Our results provide a mechanistic framework for interpreting resistance-driven outcomes in phage–bacteria systems and highlight how mathematical structure can constrain therapeutic strategies, even in simple experimental settings.

## 1 Introduction

The emergence of multidrug-resistant (MDR) bacteria is a mounting global health emergency, threatening the effectiveness of standard antibiotics in facing bacterial infections. According to the World Health Organization, antibiotic resistance is among the top 10 global public health threats. In 2021, 4.71 million (95% with uncertainty interval (UI) 4.23–5.19) deaths were associated with antimicrobial-resistant bacteria, including 1.14 million (95% UI 1.00–1.28) deaths attributable to bacterial resistance [1]. The problem is particularly acute in hospital settings, where MDR strains of, e.g., *Klebsiella pneumoniae, Acinetobacter baumannii, Pseudomonas aeruginosa*, and *Escherichia coli* are responsible for a growing number of healthcare-associated infections. These pathogens contribute, among others, to extended hospital stays, increased treatment costs, and higher morbidity and mortality [34]. Moreover, the pipeline for new antibiotics remains critically underfunded and insufficient, with few novel agents addressing the most resistant Gram-negative bacteria [17]. This confluence of rising resistance and declining innovation underscores the urgent need for alternative antimicrobial strategies.

Bacteriophages, or phages—viruses that infect and lyse bacteria—are being revisited as a viable therapeutic option in the post-antibiotic era [40]. Unlike broad-spectrum antibiotics, phages are able to amplify and spread locally, targeting microbial pathogens while also being highly specific to their bacterial hosts, minimizing off-target effects on the microbiota [35]. Phage therapy (see Ref. [21] for a recent review), first proposed nearly a century ago, was soon overshadowed by the success of antibiotics. However, the rise of MDR bacteria due to the misuse and overuse of antibiotics is renewing interest in phage therapy as a viable strategy to combat pathogenic bacterial infections. Advances in high-throughput sequencing, metagenomics, genetic engineering, and synthetic biology are now overcoming previous limitations, spurring increased support from funding agencies, biotech startups, and pharmaceutical companies. Recent studies and clinical trials have demonstrated the effectiveness of phage therapy in treating antibiotic-resistant infections, including chronic wound infections, sepsis, and respiratory tract infections caused by *Pseudomonas, Staphylococcus*, and *Acinetobacter* species [39, 6, 10, 11, 13, 19]. The adaptability of phages, their natural presence in diverse ecosystems, and their potential for genetic engineering also make them attractive tools for precision medicine and infection control. Phage cocktails and engineered phages are already being explored to broaden host range and prevent the rapid emergence of bacterial resistance during therapy [34, 26].

Effective implementations of phage therapy require a deep understanding of phage-bacteria-host population and co-evolutionary dynamics, and this is where mathematical modeling becomes useful. Models help predict the ecological and evolutionary interactions between phages and bacteria within the host, allowing for better-informed treatment designs [26]. These models can simulate the timing, dosage, and combination strategies needed for phage administration and assess synergies with antibiotics or host immune responses. Furthermore, modeling is crucial for anticipating resistance development and for designing robust phage cocktails tailored to individual infections [34]. As precision medicine continues to evolve, integrating mathematical frameworks into phage therapy protocols will be key to optimizing outcomes and mitigating the global threat of MDR bacteria.

Quantitative mathematical approaches are essential for understanding the complex dynamics of phage-bacteria interactions, particularly in simplified systems. In this context, *in vitro* experiments with MDR strains and phages provide a controlled setting to dissect key interactions and quantify the rates at which key processes occur. By means of mathematical modeling, dynamical systems theory, and statistical methods, researchers can estimate key parameters that control the population dynamics from these experimental data, such as infection rates, burst sizes, or bacterial growth [36]. These quantitative insights can enable more precise predictions of system behavior under various conditions, including different parameter values and initial conditions (i.e., monostable vs. bistable dynamics), shed light on the design of effective phage therapies, and enhance our ability to manipulate microbial ecosystems.

In this direction, a recent work by Rodríguez et al. [36] introduced a mathematical model to analyze the *in vitro* dynamics between *K. pneumoniae* and the lytic phage vB Kpn 2-P4. Through bacteria growth experiments and mathematical modeling, the authors estimated key population parameters, including bacterial growth rates, phage infection rates, burst sizes, and phage-resistance emergence probabilities. Logistic and nonlinear models were fitted using both Levenberg-Marquardt and Bayesian inference methods, yielding consistent estimates. Results showed that resistant mutants emerged rapidly and grew at rates comparable to the wild-type strain, with minimal fitness cost. These findings highlight the critical role of resistance in shaping phage therapy outcomes and the importance of integrating experimental data with quantitative models to predict and optimize potential therapeutic strategies.

In this manuscript, we analyze the qualitative dynamics of the model used in the experimental setting described in Ref. [36] (see also Fig. 1). We study the equilibria and the bifurcations, showing a coexistence scenario between phage-resistant bacteria and phages governed by a quasineutral line of equilibria (i.e., a degenerate normally hyperbolic invariant manifold, see Refs. [28, 38, 18] for other examples of quasineutral manifolds in biological models). Clearance of resistant bacteria is achievable by combining phage treatment with increased degradation of resistant strains, leading to a global transcritical bifurcation. These results highlight the potential of combined phage-antibiotic therapies to clear phage-resistant bacterial populations. Despite extensive experimental and modeling work on phage–bacteria systems, the qualitative dynamical structures governing long-term coexistence and clearance remain poorly characterized. In particular, little attention has been paid to the role of non-hyperbolic invariant sets—such as lines of equilibria—in shaping resistance-driven outcomes. Here, we address this gap by providing a rigorous dynamical systems analysis of a data-calibrated phage–bacteria model, revealing how quasineutral manifolds organize coexistence, sensitivity to initial conditions, and clearance transitions.

**Figure 1.**
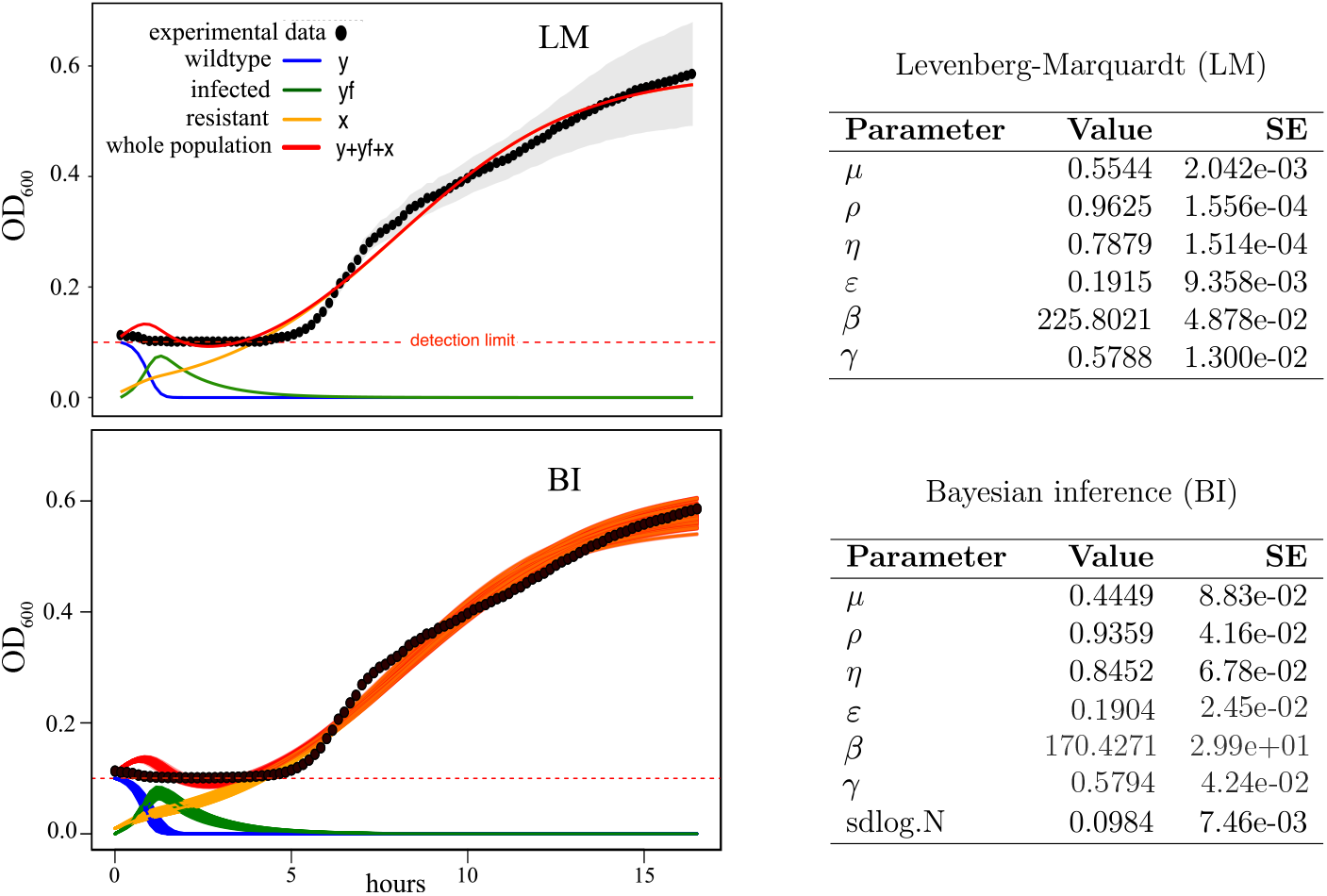
Model fit to the experimental data and parameter estimation. (Left) Dynamics of wild-type (wt) *Klebsiella pneumoniae* in the presence of the bacteriophage vB Kpn 2-P4 monitored *in vitro* (black dots). Bacterial densities are measured with the optical density (OD_600_). The dynamics obtained with the mathematical model given by Eqs. (8)-(11) and fit to the data are shown: wt population (*y*, blue lines); infected bacteria (*y*_*f*_, green lines); resistant bacteria (*x*, orange lines). The sum of all the bacterial populations (red lines) was used to compare with the experimental data. Notice that the orange lines appeared overlapped with the red ones when the phage-resistant bacteria out-competed the other strains. Fits were done with the mean populations from the experiments averaged over 21 independent replicas using two methods: the Levenberg-Marquardt (LM) and Bayesian inference (BI). (Right) Estimated parameter values using the LM and BI, with an intrinsic growth rate of the wt bacteria of *k* = 0.538 and with carrying capacity (expressed as OD) *C* = 0.887. These two latter values were obtained for *K. pneumoniae* dynamics alone (see Ref. [36] for details). See Table 1 for a detailed description of the model parameters.

## 2 Summary of the experimental results

The selected strain of *Klebsiella pneumoniae* was Kpn63, a clinical strain that belongs to the capsular type KL-64 [17]. This strain harbors a New Delhi metallo-beta-lactamase-1 (NDM-1) enzyme that confers resistance to all beta-lactam antibiotics, including carbapenems. The growth of Kpn63 was monitored without and with the presence of phage vB Kpn 2-P4 [36]. This is a lytic phage of *K. pneumoniae* isolated from sewage water [17]. The phage vB Kpn 2-P4 was selected for our study due to its broadest host range in clinical strains of *K. pneumoniae* with capsular type KL-64.

**Table 1:** Parameters of Eqs. (8)-(11). We display the symbols, the biological meaning, the values, and the units of each parameter. The values are set following the experimental results in Ref. [36]. In all these experiments *ε < γ* and *β* ≫ 1, so we assume the lower bound *β >* 1 to hold, since infected bacteria produce large amounts of phages [36]. Here CFU stands for colony-forming units and PFU for plaque-forming units. The time unit used throughout this work is [*t*] = *hours*.

| Parameter | Biological meaning | Value | Units |
| --- | --- | --- | --- |
| $k$ | Intrinsic growth rate of susceptible wild-type bacteria | $0 < k < 1$ | $t^{-1}$ |
| $\gamma$ | Replication rate of phage-resistant bacteria | $0 < \gamma < 1$ | $t^{-1}$ |
| $\mu$ | Fraction of resistant bacteria generated from susceptible ones | $0 < \mu < 1$ | Dimensionless |
| $\rho$ | Infection rate of susceptible bacteria by phages | $0 < \rho < 1$ | $\text{mL} \cdot \text{PFU}^{-1} \cdot t^{-1}$ |
| $\eta$ | Death rate of infected bacteria due to phages | $0 < \eta < 1$ | $t^{-1}$ |
| $\beta$ | Burst size (number of phages released upon bacterial lysis) | $\beta > 1$ | $\text{PFU} \cdot \text{CFU}^{-1} \cdot t^{-1}$ |
| $\varepsilon$ | Decay rate of phage-resistant bacteria | $0 < \varepsilon < 1$ | $t^{-1}$ |

The starting aliquot of the phage vB Kpn 2-P4 was stored at −80°C in SM buffer and then amplified in Luria-Bertani media supplemented with CaCl_2_ in the same bacterial strain in which it was isolated to obtain high-titer lysates. Negative (only LB media) and positive (Kpn63 without phage) controls were used to check the efficacy of phage infection. The total volume was passed through a 0.22 µm filtration unit and concentrated using CP Select InnovaPrep equipment. To determine the number of phage particles in our concentrated aliquots (phage titration), we followed the double-layer agar method. We mixed 10 µL of phage serial dilutions and 100 µL of fresh Kpn63 bacterial cultures in 3.5 mL of liquid top agar at 55°C and plated after vortexing in LB agar plates. After overnight incubation at 37°C, plaque-forming units (PFUs) were counted, reaching a final titer of 1.5 × 10^10^ PFU mL^−1^. The appearance of the plaques was small and cloudy; the turbidity of the halos can result from partial bacterial lysis or the activity of phage-associated enzymes (depolymerase), which degrade components of the bacterial cell wall or extracellular matrix without lysing the bacteria.

To study the infectivity of the phage and the phage-bacteria dynamics in liquid medium, we determined the strength of lysis based on time-lapse turbidity (OD_600_) measurements at 10-min intervals. Our experiments were performed in 96-well plates, incubated at 37°C inside a plate reader (Multiskan) for 16.5 hours. Once we knew the correlation of CFU mL^−1^ from our bacterial suspensions and PFU mL^−1^ from the phage aliquots, we tested different initial multiplicity of infection (MOI) conditions by combining 150 µL of bacterial suspension and 10 µL of phage dilution in each well. All experiments were performed with 24 independent replicates per condition (MOI 1, 0.1, 0.01, and 0.001).

## 3. Mathematical model

In this section we introduce the mathematical model describing the bacteria-phage dynamics used in the experiments carried out in Ref. [36]. Since the experiments were done in a liquid medium, a mean field model based on ordinary differential equations (ODEs), i.e., the limit of infinite diffusion, is used. The state variables of the model are the density of wild-type (wt) *Klebsiella* strains *y*(*t*), infected *Klebsiella* strains *y*_*f*_ (*t*), phages *ϕ*(*t*), and phage-resistant *Klebsiella* strains *x*(*t*). The variable *x* may be either considered as a single type of phage-resistant mutant or as an average population of different resistant mutants. The key modelled processes, shown in Fig. 2(b), can be represented by the following stoichiometric relations:

**Figure 2.**
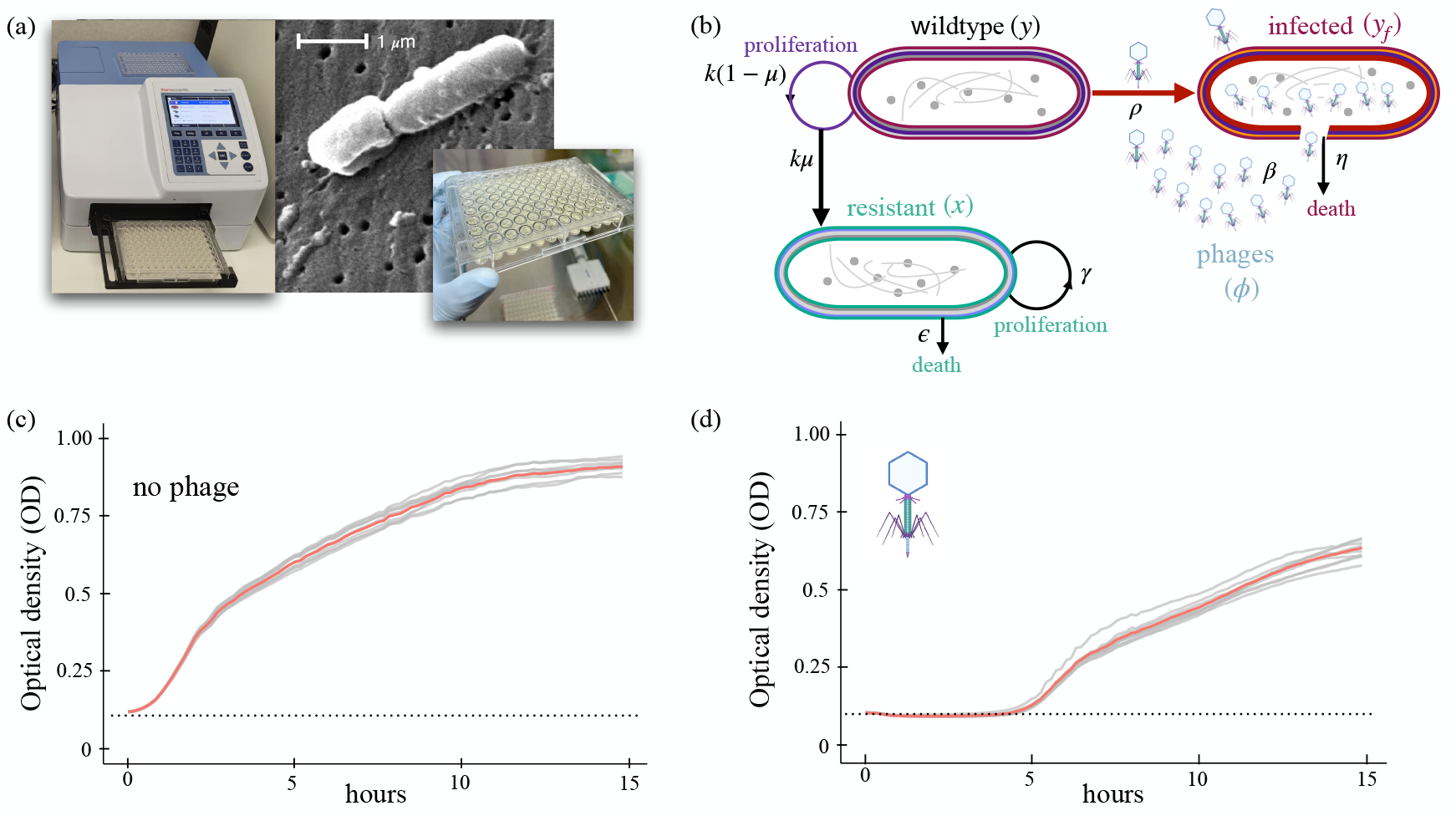
Experimental system and modeled interactions. (a) Experimental setup to track the population dynamics of *Klebsiella pneumoniae* (strain Kpn63). The electron microscope image shows two bacteria cells (image by Janice Carr/CDC—CDC, Atlanta (USA) from Wikipedia). (b) Processes modeled in this work. We consider a wild-type (wt) population of bacteria (*y*) that can become infected (*y*_*f*_) by phages (*ϕ*). The wt strains can mutate during replication and generate phage-resistant cells (*x*). The stoichiometry of the whole interactions is provided in reactions (1)-(7), modeled with Eqs. (8)-(11). (c) Growth curves for *Klebsiella pneumoniae* (strain Kpn63) without phages measured with the optical density (OD). We show eight replicates with an initial condition of bacteria with OD = 0.1. (d) Kpn63 dynamics with an initial population also with OD = 0.1 in the presence of phages. The horizontal dotted lines show the limit detection of the OD measures.

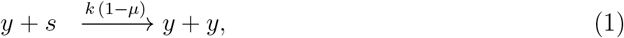

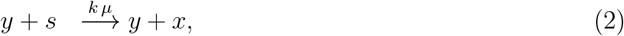

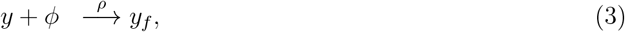

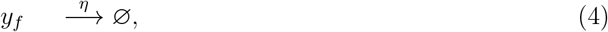

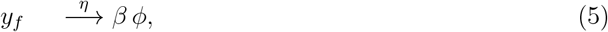

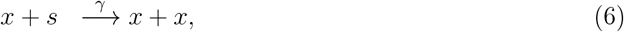

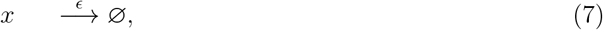

Reaction ((1)) is the duplication of *Klebsiella* wt cells, which depends on available nutrients (*s*, not explicitly considered) and is proportional to the intrinsic growth rate *k >* 0, with *µ* ∈ [0, 1] being the probability of producing a resistant strain. Notice we assume in reaction (2) that resistant strains arise due to mutations during wt replication. Reaction ((3)) is the infection of wt cells, which is proportional to the infection rate *ρ >* 0. Infected cells die because they are lysed by the phages at a rate *η >* 0, see reaction ((4)). When these cells are lysed, they release new phages at a rate *η* [reaction (5)], being *β* ≫ 1 the burst size (number of viral particles released upon lysis). Phage-resistant cells duplicate at a rate *γ >* 0 [reaction (6)] and they die at a rate *ϵ >* 0 [reaction (7)]. The above set of reactions can be modeled using the law of mass action, giving rise to a dynamical system governed by the following set of ODEs:

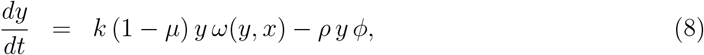

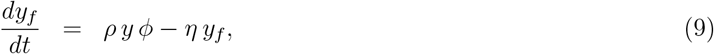

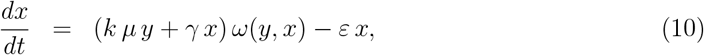

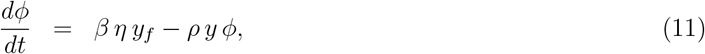

where the function

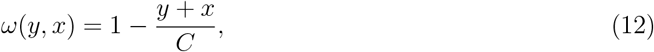

is a logistic growth constraint including competition between the bacterial strains that duplicate.

Several modeling assumptions deserve comment. We neglect superinfection of already infected cells and explicit phage decay, assumptions justified by the short experimental time scales and the large burst sizes observed in vitro. The decay term for phage-resistant bacteria represents an effective mortality that may arise from fitness costs, immune-mediated clearance, or synergistic antibiotic effects. Importantly, the qualitative dynamics reported here do not depend on the specific biological origin of this term, but rather on its magnitude relative to the growth rate of the resistant population.

The model further assumes that infected bacteria do not proliferate and therefore do not compete for nutrients; accordingly, this population is excluded from the logistic growth term. Although the proliferation rate of phage-resistant bacteria is allowed to differ from that of the wild-type strain, experimental measurements indicate that this difference is small [36]. Once a wild-type cell is infected, it is assumed to be refractory to further infection. All model parameters, together with their biological interpretation, numerical values, and units, are summarized in Table 1 (see also Fig. 1).

## 4. Results and discussion

In this section, we first compute the equilibria of the system (8)-(11). Second, we study their stability and bifurcations. Finally, we provide a sensitivity analysis of the system response upon changes in parameters and initial conditions by exploring the variational equations.

### 4.1. Equilibria and stability analysis

The equilibrium points of the system (8)-(11) are those points 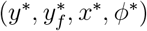 satisfying

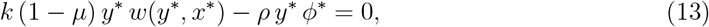

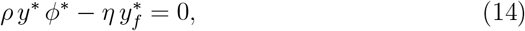

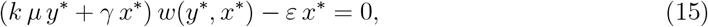

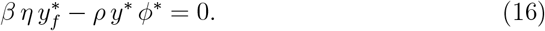

We calculate the solutions of this system, distinguishing two cases: (I) *y*^∗^≠ 0 and (II) *y*^∗^ = 0.

**Case I:** *y*^∗^≠ 0. From the condition (14), it follows that 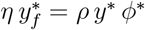. Substituting in (16), we obtain (*β* − 1) *ρ y*^∗^ *ϕ*^∗^ = 0. Since *β >* 1 and *ρ >* 0, we conclude that *ϕ*^∗^ = 0 and, therefore, 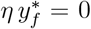. Since *η >* 0, we also obtain 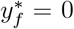. Under these conditions, the equation (13) reduces to *w*(*y*^∗^, *x*^∗^) = 0. Then, *y*^∗^ = *C* − *x*^∗^. Substituting in (15), it follows that *x*^∗^ = 0 because *ε >* 0. Hence, we obtain an equilibrium point at (*C*, 0, 0, 0), which we denote hereafter by *P*_1_. This equilibrium point involves the persistence of only the *Klebsiella pneumoniae* wt strain achieving the maximum population (i.e., the carrying capacity *C*).

**Case II:** *y*^∗^ = 0. First, note that the condition (13) is trivial in this case. From the condition (14), it follows that 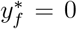. Under these conditions, the equation (16) is satisfied, and (15) is equivalent to the equation

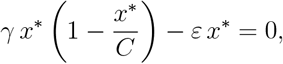

for any value of *ϕ*^∗^. Therefore, we obtain two lines of equilibria, determined by the two solutions of the above expression, *x*^∗^ = 0 and *x*^∗^ = *C*(1 − *ε/γ*). In what follows, we denote these two straight lines of equilibria by

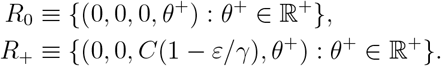

#### Remark 1

We take only the biologically meaningful part of the straight lines, *θ*^+^ ≥ 0. In addition, we also require the condition *C*(1 − *ε/γ*) *>* 0 to be satisfied, i.e., *ε < γ*, in agreement with the experiments performed in Ref. [36].

Before presenting the formal stability results, it is useful to outline the structure of the phase space at a conceptual level. The system admits equilibria corresponding to wild-type dominance, phage extinction, and resistant–phage coexistence. Remarkably, the latter does not occur at an isolated point but along a continuum of states parameterized by phage density. As shown below, the stability properties transverse to this continuum determine whether the system converges to coexistence or clearance. The following result follows.

#### Theorem 1

*System* (8)*-*(11), *with parameter values restricted to those given in Table 1, has an unstable hyperbolic equilibrium point at P*_1_ = (*C*, 0, 0, 0), *which is of saddle type (dim*(*W* ^*s*^(*P*_1_)) = 3 *and dim*(*W* ^*u*^(*P*_1_)) = 1*). Moreover, under the condition ε < γ, the system has two non-negative half-lines of equilibria, which satisfy*

*i*) *R*_0_ = {(0, 0, 0, *θ*^+^) : *θ*^+^ ∈ ℝ^+^} *is unstable*,

*ii*) *R*_+_ = {(0, 0, *C*(1 − *ε/γ*), *θ*^+^) : *θ*^+^ ∈ ℝ^+^} *is unstable for* 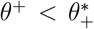 *and locally stable for* 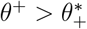, *where* 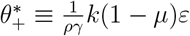.

*Proof*. The Jacobian matrix of the system (8)-(11) is given by

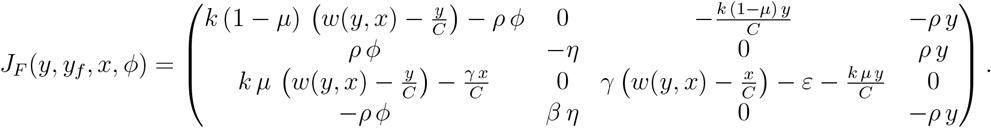

The characteristic polynomial associated with the Jacobian matrix at the equilibrium point *P*_1_ is

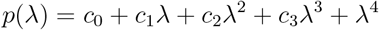

with

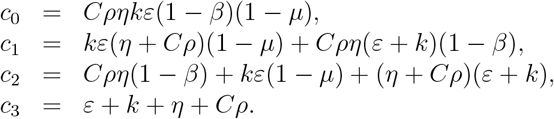

By carefully manipulating the above polynomial expression, we write *p* as the product of the following two quadratic polynomials

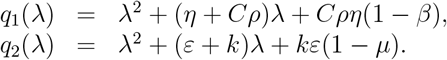

Therefore, the eigenvalues related to the equilibrium point *P*_1_ are easily computed as the roots of *q*_1_ and *q*_2_. First, we compute the roots of *q*_1_,

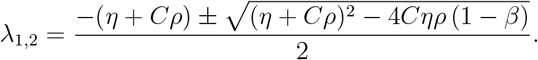

Since the parameters of our model take positive values and *β >* 1, the discriminant is always positive, Δ = (*η* + *Cρ*)^2^ + 4*Cηρ* (*β* − 1) *>* 0. Moreover, as 4*Cηρ* (*β* − 1) *>* 0, it follows that 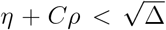. Hence, these eigenvalues are real and of opposite sign, *λ*_1_ *>* 0 and *λ*_2_ *<* 0.

Second, the roots of *q*_2_ are

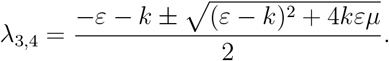

We obtain that both eigenvalues are real, since (*ε* − *k*)^2^ + 4*kεµ >* 0. Regarding the sign of the roots, it is clear that *λ*_4_ *<* 0. As for the sign of *λ*_3_, since 4*kε*(1 − *µ*) *>* 0, it follows that

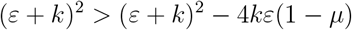

and

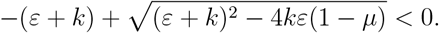

Therefore, this eigenvalue is negative (*λ*_3_ *<* 0). In summary, *P*_1_ is an unstable hyperbolic equilibrium point of saddle type, with dim(*W* ^*s*^(*P*_1_)) = 3 and dim(*W* ^*u*^(*P*_1_)) = 1.

1. The Jacobian matrix associated with the line of equilibria *R*_0_ = (0, 0, 0, *θ*^+^) : {*θ*^+^ ≥ 0} is given by the block-lower-triangular matrix

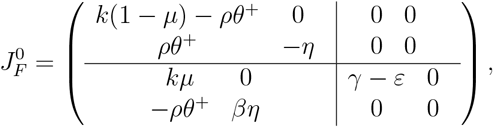

whose spectrum is directly computed:

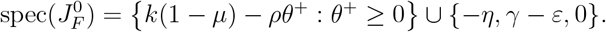

As *γ > ε*, all equilibrium points on *R*_0_ have an unstable direction (associated with the positive eigenvalue *γ* − *ε*). Therefore, the line of equilibrium points *R*_0_ is unstable.
2. The Jacobian matrix *J*_*F*_ evaluated at points on the half-line *R*_+_ = {(0, 0, *C*(1−*ε/γ*), *θ*^+^) : *θ*^+^ ∈ ℝ^+^} is given by the block-lower-triangular matrix

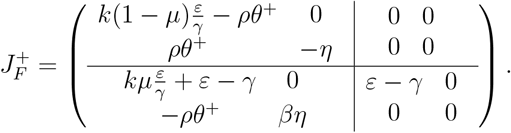

We easily obtain

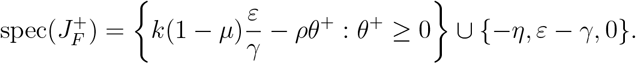

Depending on the values of *θ*^+^, we distinguish two cases. On the one hand, there exists an unstable direction when 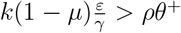. By defining 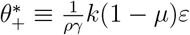, we obtain the line

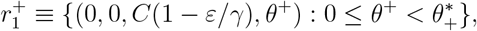

which consists of unstable equilibrium points. On the other hand, we get the line

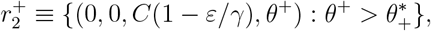

consisting of equilibrium points with three stable directions and one central direction.

#### Remark 2

The *ϕ*-direction related to the zero eigenvalue, which appears respectively in the spectrum of 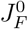 and 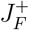, is entirely linked to the existence of the half-lines of equilibrium points. Each half-line, *R*_0_ and *R*_+_, acts as a degenerate normally hyperbolic invariant manifold. Therefore, its dynamics are governed by the transversal hyperbolic directions.

To complete the analysis carried out in this section, we consider the dynamics of the system by using the values of the parameters estimated with the Bayesian inference (BI) in the experiments [36] (see Fig. 1). We notice that using the values obtained with the Levenberg-Marquardt (LM) method would result in the same qualitative dynamics. The equilibrium points using these estimated values are given by

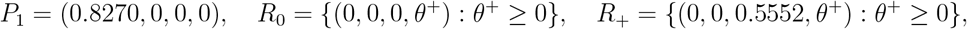

and the spectra of their associated Jacobian matrices are

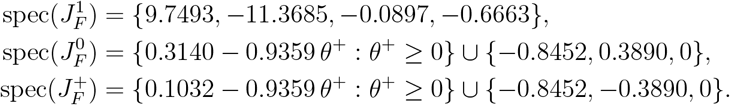

Following the results above, considering the estimated parameter values from the data, the orbits will move towards the half-line *R*_+_, since the other equilibria, the point *P*_1_ and the half-line *R*_0_, are unstable. Due to the transverse stability of 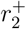, the only asymptotic state the system can achieve is the stable part of the half-line *R*_+_. The point separating the stable and unstable regions along the line *R*_+_ is given by 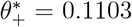, with *x*^∗^ = 0.5552. This prediction is in agreement with the values obtained for the phage-resistant mutants in the experiments. The fits performed with both the LM and the BI showed a population of phage-resistant cells rapidly out-competing the susceptible ones together with the extinction of the infected bacteria. The dynamics for the phage-resistant cells started saturating at approximately 15-16 hours, achieving a population value of *x*(*t* = 16) ≈ 0.56 (see Fig. 1).

### 4.2. Bifurcations

The computation of the equilibria of Eqs. (8)-(11) and the analysis of their linear stability allow us to identify the parameters that cause qualitative changes in the dynamics. That is, the identification of bifurcations. Most of these bifurcations, with one exception, are not biologically relevant, as they occur for parameter values outside the biologically admissible range.

Among all the assumptions made regarding the model parameters (see Table 1), there is only one that can be modified in our model without entirely losing biological meaning: *γ > ε*. Although such a bifurcation will never occur in the case of the bacteria strain studied here, that is, under the estimated parameter values, both the parameters *γ* and *ε* could vary for other bacterial strains. Slowing the growth of multidrug-resistant (MDR) bacteria that have also developed resistance to bacteriophages is highly challenging and likely requires multifaceted strategies. One approach could involve phage cocktails or engineered phages, which target multiple bacterial receptors or deliver lethal genetic payloads [21]. CRISPR-Cas systems, delivered via phages or conjugative plasmids, can precisely eliminate resistant strains by targeting essential or resistance genes [8]. Bacteria often pay a fitness cost for resistance, which could be exploited by adjusting environmental conditions (e.g., nutrient limitation) or by introducing competitive strains weakening the growth of the phage-resistant strains. Quorum sensing inhibitors can also disarm bacteria without selecting for further resistance [32]. Moreover, phage-resistant bacteria could slow down growth or increase mortality if an antibiotic acting synergistically with bacteriophages is introduced. In this sense, phage-antibiotic therapies have shown promise in resensitizing bacteria to treatments [9].

In what follows, we analyze the bifurcation associated with the extinction of phage-resistant bacteria. All parameters are fixed at the values estimated using the Bayesian inference method (Fig. 1), with the exception of *ε*, which is treated as the bifurcation parameter. From a biological and therapeutic standpoint, bifurcations in this system correspond to qualitative shifts in treatment outcomes. Among all parameters, the effective mortality rate of phage-resistant bacteria (*ε*) plays a distinctive role, as it can be experimentally modulated through antibiotic exposure, immune-mediated clearance, or engineered phage strategies. We therefore examine *ε* as a control parameter governing transitions between resistant persistence and clearance.

First, we note that the spectrum of the Jacobian matrix at *P*_1_ varies with *ε*, yet the instability of *P*_1_ remains unaffected by the condition *γ > ε*. Accordingly, we focus solely on the changes along the half-lines of equilibria *R*_0_ and *R*_+_ as the bifurcation parameter varies. It is clear that the condition *ε* = *γ* marks a change in the stability of the equilibrium lines in the *x*-direction, as suggested by the spectrum of the Jacobian associated with both lines. To explore this stability change in more depth, we restrict our analysis to the invariant plane Π = {*y* = 0, *y*_*f*_ = 0}. Since 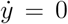 and 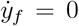 under this restriction, the 4-dimensional system (8)-(11) is reduced to a 2-dimensional model. Moreover, on the invariant plane Π, we also obtain that 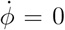 for all values of *ϕ*. Thus, within this invariant plane, the variable *ϕ* remains constant and can be treated as a parameter in the reduced system. This means that the dynamics are effectively governed by the below differential equation:

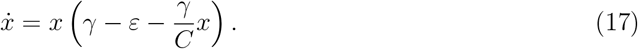

After the change of variable 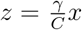, Eq. (17) can be written in the normal form of the transcritical bifurcation, i.e., *z*? = *αz* − *z*^2^ with *α* = *γ* − *ε*. Fig. 3 illustrates the dynamics of *x* and *ϕ* on the plane Π, by means of phase portraits for various values of the bifurcation parameter.

**Figure 3.**
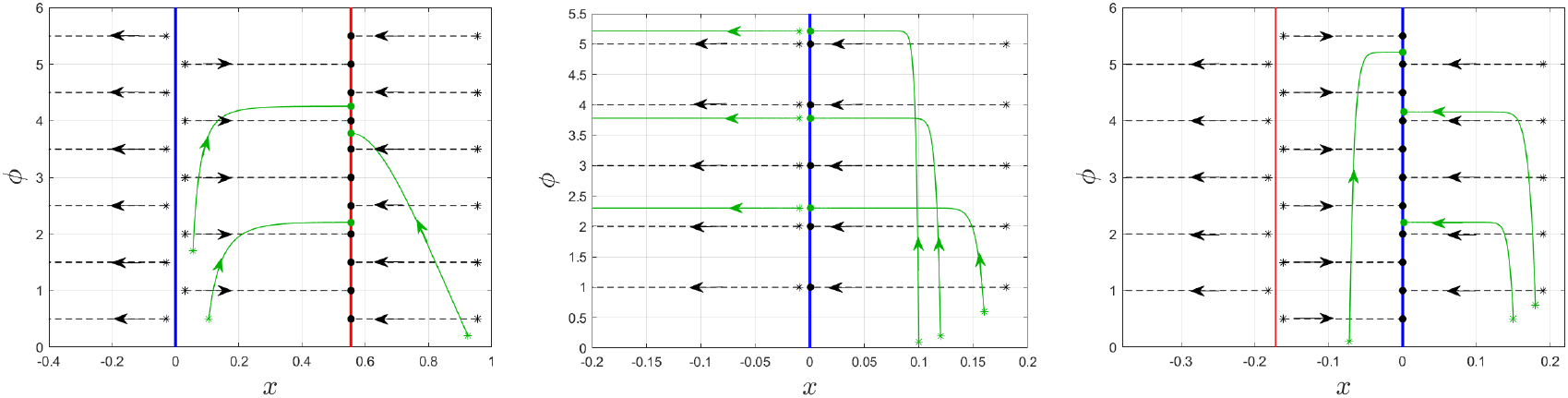
Phase portraits and bifurcations. Orbits of the system (8)–(11) projected on the invariant plane Π = {*y* = 0, *y*_*f*_ = 0} for different values of *ε*. The other parameters are fixed, as indicated in the table of Fig. 1 (estimated with the Bayesian inference (BI)). The half-lines of equilibrium points *R*_+_ and *R*_0_ are displayed in red and blue, respectively. Initial conditions are marked with asterisks, and the end of orbits with dots. Orbits with initial conditions on the plane Π are shown in black dashed lines, while those with initial conditions *y* = 0 and *y*_*f*_≠ 0 (projected onto the invariant plane) are shown in green lines. (Left) *ε < γ* with *ε* = 0.1904: The line *R*_+_ is an attractor, and the line *R*_0_ is repulsive. (Middle) *ε* = *γ* with *ε* = 0.5794: Both half-lines coincide (purple), forming a single line of saddle-node equilibrium points. (Right) *ε > γ* with *ε* = 0.7: The line *R*_+_ becomes unstable and negative on the *x*-axis, losing biological meaning, while the line *R*_0_ becomes stable. Orbits shown for negative values of *x* correspond to cases with no biological meaning.

Although the transcritical bifurcation in the *x*-direction is fully characterized, it is important to clarify what occurs in the other two directions that were not considered. In fact, the eigenvalue in the *y*_*f*_-direction, as verified in the previous section, is always negative (−*η <* 0). Therefore, the only direction that may cause instabilities is the *y*-direction. We describe the three cases to determine whether the dynamics of the equilibrium points along the half-lines indeed change.

**Case I:** *ε < γ*. This case corresponds to the experimental data reported in Ref. [36]. In the left panel of Fig. 3, the two half-lines of equilibrium points *R*_0_ and *R*_+_ are shown. Indeed, the half-line *R*_0_ (blue) is unstable while the half-line *R*_+_ (red) is stable, on the invariant plane Π. Based on the stability analysis above, see Theorem 1, we proof that when *y*≠ 0 and *y*_*f*_≠ 0, the half-line *R*_+_ splits into two pieces with different stability properties: 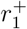 unstable and 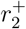 stable. We conclude that 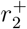 is an attracting half-line of equilibrium points (see Fig. 4, left panels).

**Figure 4.**
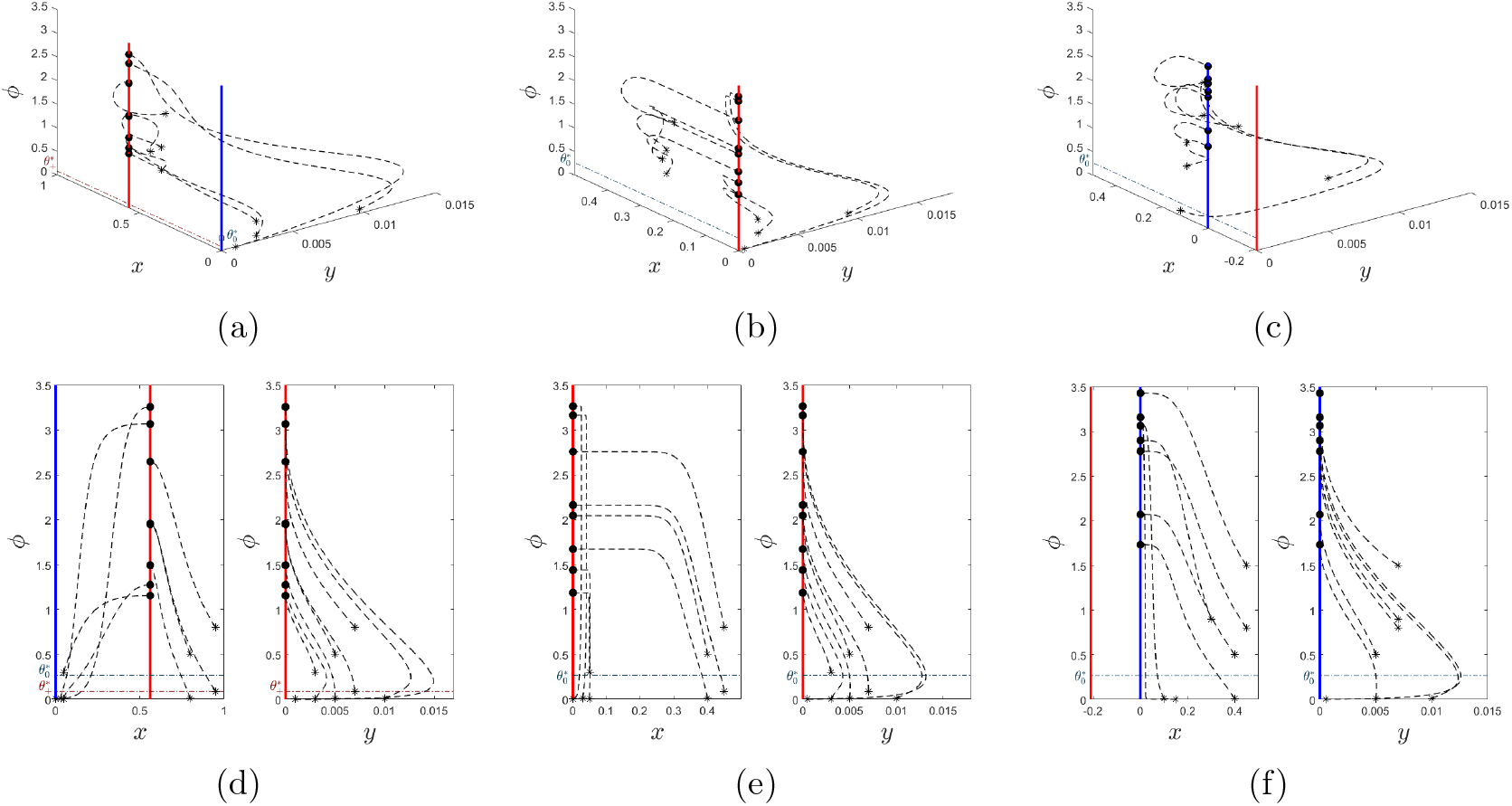
Orbits of the system (8)–(9) around the bifurcation associated with the parameter *ε*. The line *R*_+_ is shown in red and the line *R*_0_ in blue. The start of each orbit is indicated by an asterisk and the end by a dot. For all simulations, parameter values are taken from the table in Fig. 1, except for the bifurcation parameter *ε*, which is varied. (a)–(c) Orbits of the populations *x, y*, and *ϕ* for different initial conditions outside the invariant plane {*y* = 0, *y*_*f*_ = 0}. (a) Example with *ε* = 0.1904 *< γ*. The line *R*_+_ is attractive, while the line *R*_0_ is repulsive. (b) The case *ε* = *γ* = 0.5794 makes both lines coincide, forming a single equilibrium line *R*_0_ of saddle type. (c) Here *ε* = 0.7 *> γ* is selected. The line *R*_+_ becomes unstable and loses biological relevance, while the line *R*_0_ becomes stable. (d)–(f) Projections of the orbits from panels (a)–(c), respectively, onto the (*x, ϕ*) and (*y, ϕ*) planes. We only show nonnegative orbit values with biological meaning.

**Case II:** *ε* = *γ*. Here, the two equilibrium half-lines coincide,

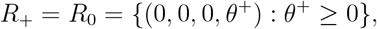

as shown in Fig. 3 (middle panel). The behavior is completely determined in all directions except the *y*-direction, which remains dependent on the value 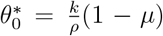. For all 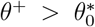, the equilibria are stable for any initial condition in the biologically feasible region (*x >* 0). When 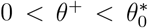, the equilibria are unstable. These dynamics are illustrated in Fig. 4, specifically in panels (*b*) and (*e*).

**Case III:** *ε > γ*. In this case, for the equilibrium half-line *R*_+_, the *x*-component is given by *x* = *C*(1 − *ε/γ*) *<* 0. Thus, it lies outside the biologically admissible region (in any case, as seen in Fig. 3, right panel, it features an unstable direction due to the eigenvalue *ε* − *γ >* 0). The stability of the points on the line *R*_0_ is completely determined in all directions except the *y*-direction, which depends on the critical value 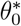. Thus, we define the half-lines:

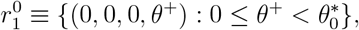

which contains unstable equilibrium points (due to the instability in the *y*-direction), and

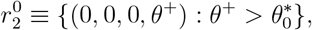

which contains stable equilibrium points; in other words, 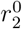 is an attractor half-line of equilibria (see Fig. 4, right panels).

As shown in the different panels of Fig. 4, for initial conditions with *ϕ* below a certain threshold, the orbits move away from the equilibrium points. However, once the orbit crosses this threshold (denoted 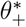 in the case *ε < γ* and 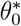 in the other cases), attraction toward the corresponding equilibrium line begins. Furthermore, we observe that the higher the distance from the attracting equilibrium line, the faster the convergence; this is reflected in the eigenvalue in the *y*-direction becoming increasingly negative. Moreover, the local stability analysis of *R*_+_, together with the behavior illustrated in Fig. 4, panels (*d*) and (*f*), allows us to propose a biologically significant interpretation of the observed phenomenon. For very small initial phage populations *ϕ*_0_, the virus appears to develop a mechanism allowing it to persist, even when the bacterial population is significantly larger. Notably, there is an initial sharp decline in the resistant bacterial population, accompanied by a rapid growth of the phage population. From a biological perspective, if experimental validation confirms this mechanism, it could be exploited to induce a drastic initial reduction in resistant bacteria.

In summary, we have identified a global transcritical bifurcation in the *x*-direction involving the two half-lines of equilibria *R*_0_ and *R*_+_. We name it a global transcritical bifurcation because the collision and interchange of stability is not between two single points, but between a continuum of points at each line. Additionally, we have shown that, in the full system, the attracting half-line 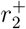 (present when *ε < γ*) loses stability in favor of 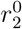 as the bifurcation parameter *ε* crosses the critical threshold *γ*.

### 4.3. Sensitivity Analysis

In this section, we provide a local sensitivity analysis of the model exploring how changes in parameters and initial conditions impact the dynamics. Our approach follows the study of the variational equations outlined in Ref. [31]. Sensitivity analysis also provides key insights into the robustness of the mathematical model when subjected to uncertainties, whether in the model parameters or in the initial conditions. In the previous section, the parameter *ε* was identified as a key factor influencing the system’s dynamics, acting as a bifurcation parameter. Given its importance, we begin the sensitivity study by examining the effects of small perturbations in *ε*. We then extend the same methodology to the remaining parameters of the model.

We denote

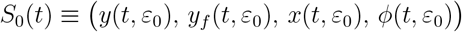

as the solution of the system (8)–(11), where *ε*_0_ is a fixed value of the parameter, and 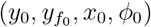 are the initial conditions.

Given any value *ε* such that |*ε* − *ε*_0_| ≪ 1, we analyze how the solution of the perturbed system

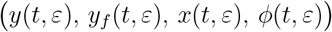

evolves relative to the reference solution *S*_0_(*t*). To this end, we consider a power series expansion of the solution with respect to *ε* in a neighborhood of *ε*_0_. For example, the function *y*(*t, ε*) can be expanded as follows:

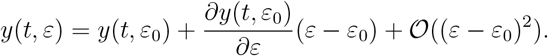

In this way, we can identify the leading-order terms that contribute to the difference between trajectories. Retaining only the first-order perturbation, we obtain

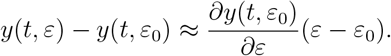

That is, a large value of the term *∂y*(*t, ε*_0_)*/∂ε* implies a greater discrepancy between the trajectories, and therefore a higher sensitivity to the parameter *ε*.

To compute these partial derivatives, we numerically approximate the solution *S*_0_(*t*) of the system (8)–(11) and substitute it into the following system of variational equations associated with the model:

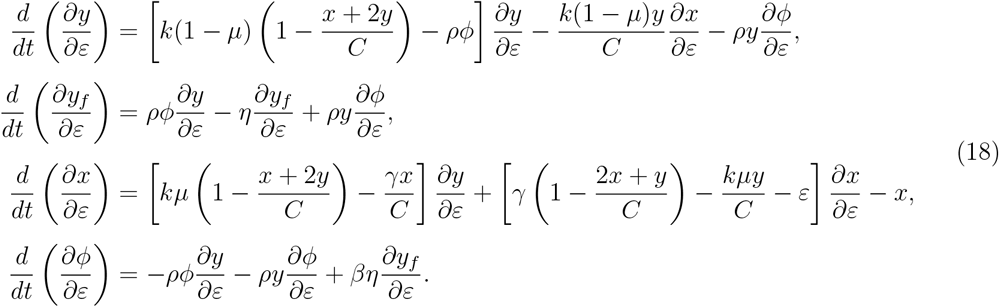

Since the original system and the perturbed one share the same initial conditions, the variational equations must reflect this fact as well. Thus, they satisfy the following initial conditions:

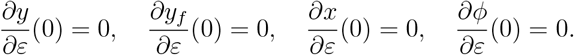

We can numerically estimate the solutions of the system (18). The results of the analysis for the parameter *ε* show that the variations for *y, y*_*f*_, or *x* are very small, being significant only for *ϕ*, as illustrated in Fig. 5. Actually, a value of *∂ϕ/∂ε* of about ∼10 is not a very significant variation considering the order of magnitude of the phage population. In principle, we only consider a fluctuation truly significant when the order of magnitude of the derivative is higher (order 2 or above),

**Figure 5.**
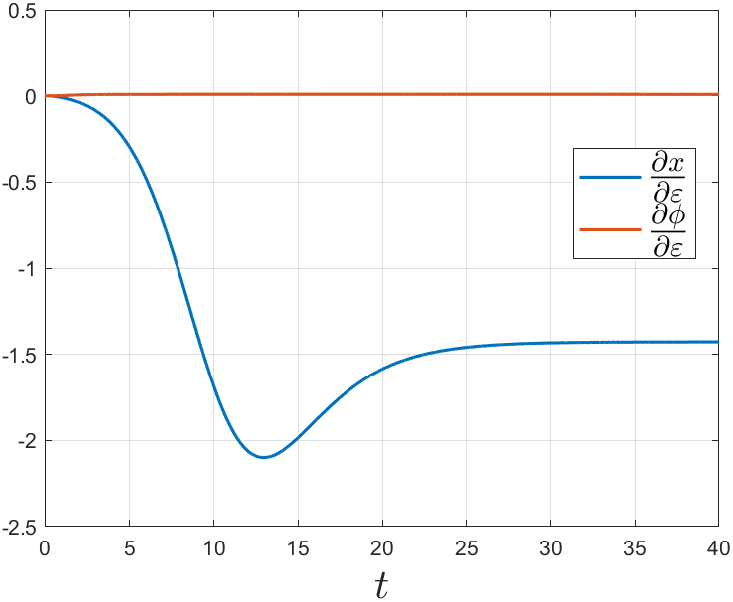
Sensitivity analysis of *x* and *ϕ* as functions of *ε* over the time interval *t* ∈ [0, 40], for the system given by equations (8)-(11). The estimated parameter values given in the table in Fig. 1 are used. Additionally, an initial condition very similar to that of the experiments [36] is taken, particularly, 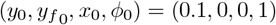. The variable *x* shows a variation that increases slightly at the beginning but quickly stabilizes. The variable *ϕ* barely shows any variation. Only *x* and *ϕ* are shown in the figure, as *y* and *y*_*f*_ are negligible.

Next, we repeat this analysis for any parameter *λ* ∈ {*β, η, γ, ν, ρ, k*} of the model. The results of the analysis show that, as in the case of the parameter *ε*, the variations for *y* or *y*_*f*_ are very small, being of interest only those for *x* and *ϕ*. We illustrate the results obtained in Fig. 6. We conclude that the model (8)-(11) is quite robust in the sense that the results are not conditioned by perturbations in the parameters. However, we should note that in the particular case of the variable *ϕ* some sensitivity to the parameters can be appreciated, particularly to *η*, where the highest peak value of all is reached (see Fig. 6, panel (b)).

**Figure 6.**
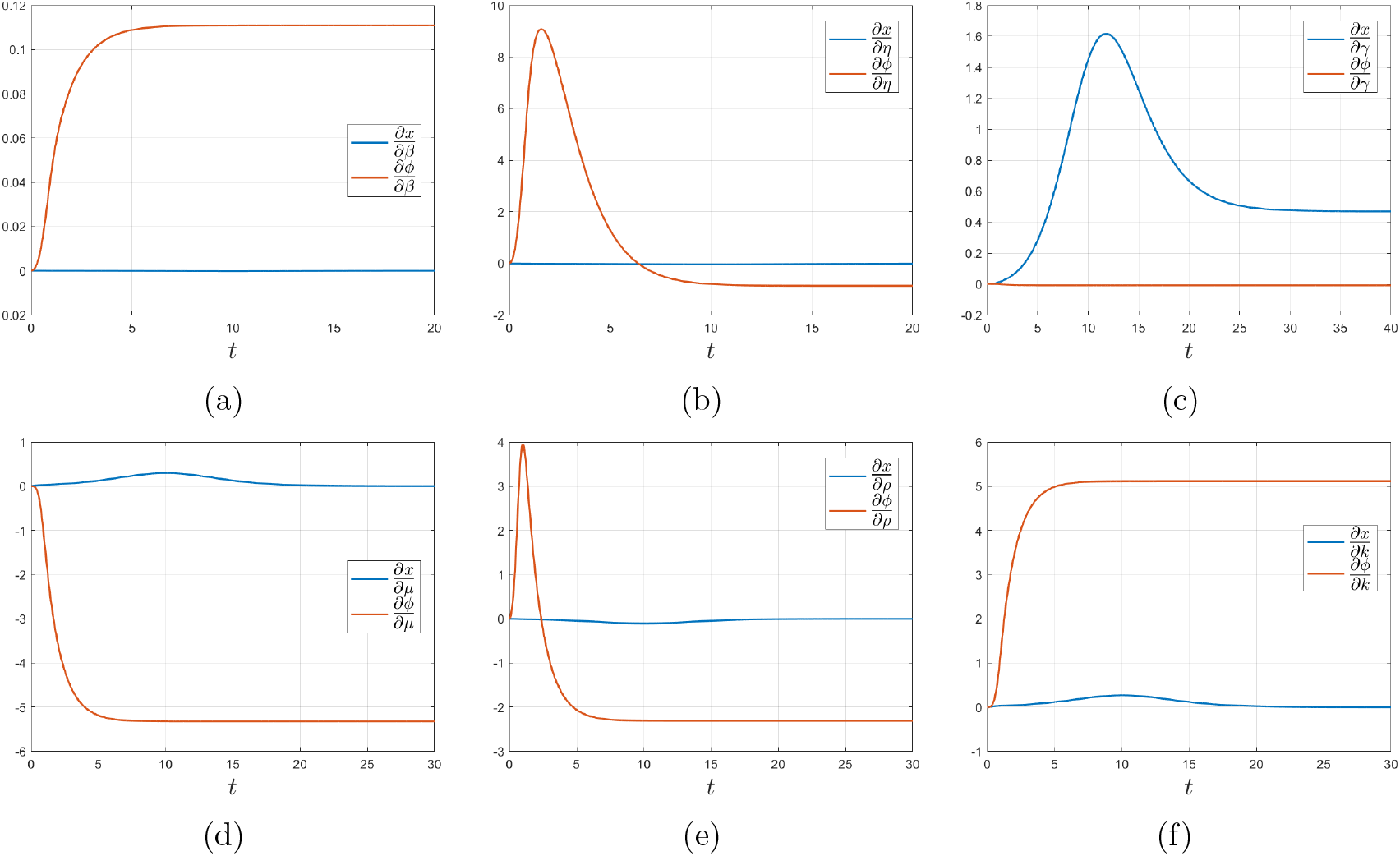
Sensitivity analysis of *x* and *ϕ* as functions of the parameters of the system (8)-(11) for different time intervals. The estimated parameter values given in the table in Fig. 1 are used. Additionally, an initial condition very similar to that of the experiments [36] is taken, that is, 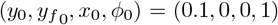. (a) The variable *ϕ* shows a small variation with respect to parameter *β*, while *x* practically shows no variation. (b) The variable *ϕ* has a notable initial variation with respect to *η*, although it rapidly decreases. On the other hand, *x* shows no variation. (c) The variable *ϕ* barely fluctuates with respect to *γ*, while the variable *x* varies slightly at the beginning. (d) The variable *ϕ* varies rapidly initially with respect to *µ*, while the variable *x* barely moves. (e) The variable *ϕ* varies slightly with respect to *ρ* at the beginning and the variable *x* barely varies. (f) The variable *ϕ* shows some variation with respect to *k*, and the variable *x* shows hardly any changes.

To complete the sensitivity analysis of the model, we study its behavior under variations in the initial conditions. Of particular interest is the sensitivity with respect to the initial phage concentration, *ϕ*_0_. To achieve this, we apply a method based on parameter sensitivity analysis, relying in particular on variational equations. We denote by

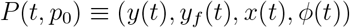

the solution of the system (8)–(11) with initial conditions

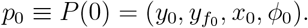

Given a small perturbation Δ ≪ 1, we study the evolution of the solution to the perturbed system:

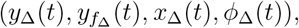

where the initial conditions are now

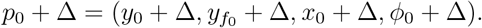

We use a Taylor expansion of each component of *p*_0_ + Δ about *p*_0_:

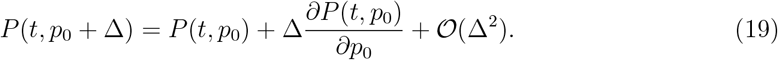

Differentiating with respect to time yields

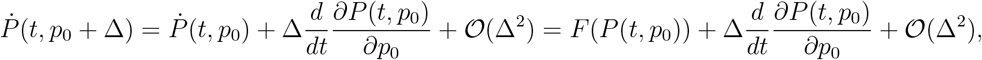

where *F* (·) denotes the vector field of the system (8)–(11). Since *P* (*t, p*_0_ + Δ) is also a solution of the system, we can expand *F* around *P* (*t, p*_0_) to obtain

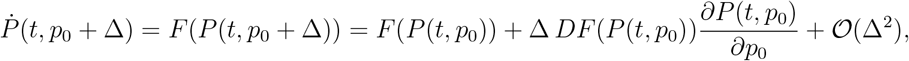

where *DF* is the Jacobian matrix of *F* evaluated at *P* (*t, p*_0_).

Equating the above expressions for 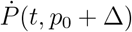 and neglecting higher order terms, we conclude that the sensitivity matrix satisfies

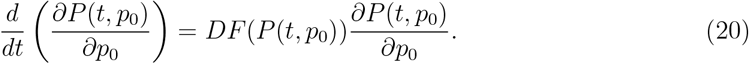

Furthermore, considering the first-order perturbation and evaluating (19) at *t* = 0, we have

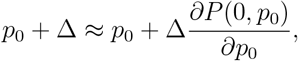

which implies the initial condition for the sensitivity matrix

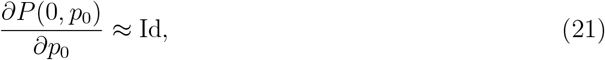

where Id is the 4 × 4 identity matrix.

The system of equations given by (20) corresponds precisely to the variational equations associated with the sensitivity to initial conditions. Writing this system in matrix form, we have

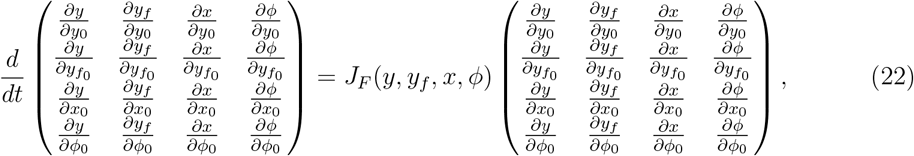

where the Jacobian matrix *J*_*F*_ is given by

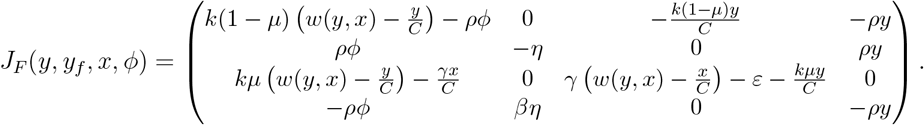

In total, this system consists of 16 coupled ordinary differential equations, with initial conditions specified by (21). We numerically approximate the solutions of system (22), and the results are illustrated in Fig. 7.

**Figure 7.**
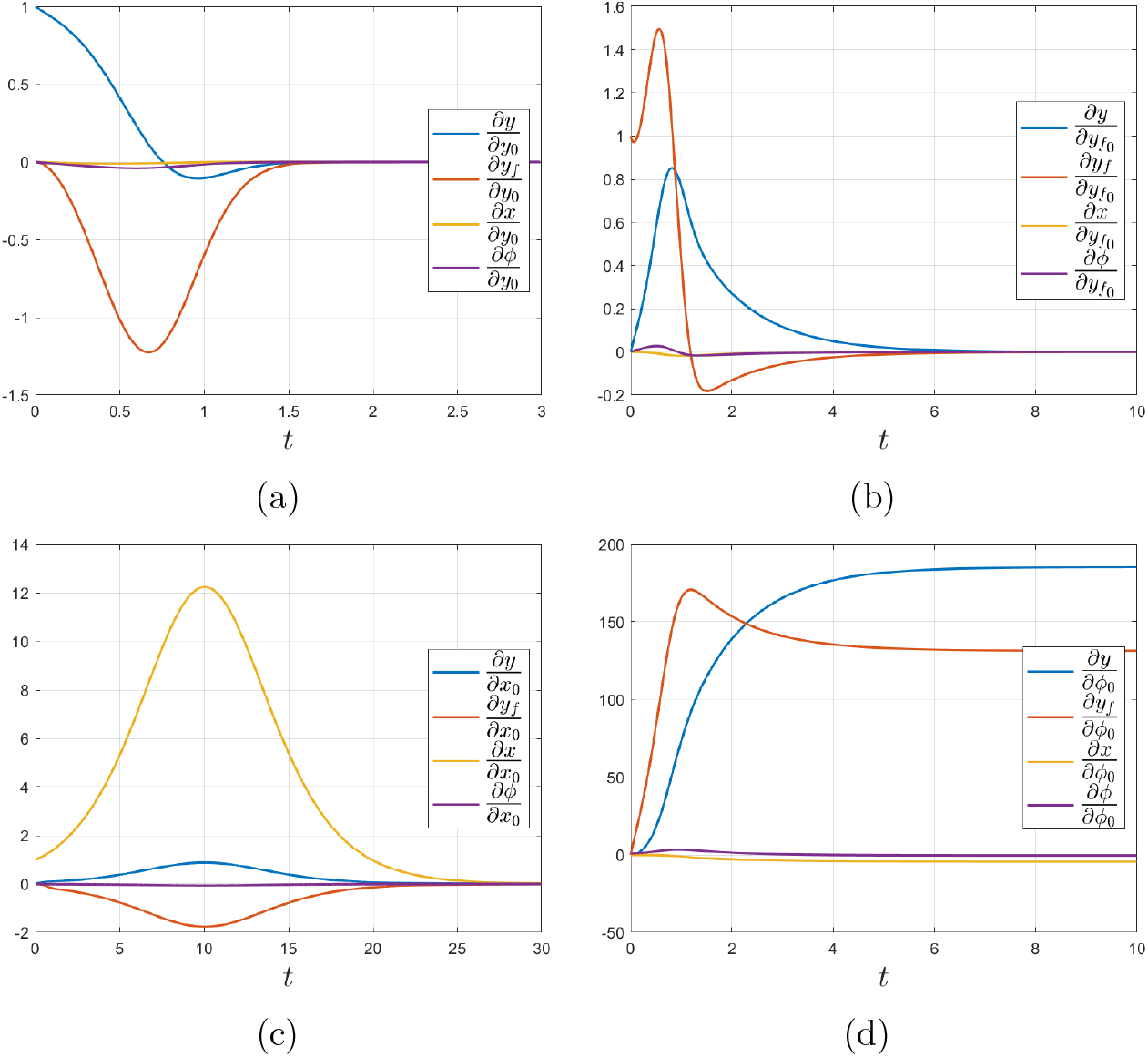
Sensitivity analysis of *y, y*_*f*_, *x*, and *ϕ* with respect to the initial conditions of the system (8)–(11) over different time intervals. The parameter values used correspond to those estimated in the table in Fig. 1. Additionally, the initial condition is chosen to closely resemble that of the experiments in Ref. [36], specifically 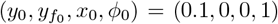. (a) The variables *y* and *y*_*f*_ exhibit a small initial variation with respect to the initial condition *y*_0_ that rapidly decreases, while the variables *x* and *ϕ* show almost no change. (b) All variables except *ϕ*, which remains nearly constant, show an initial variation relative to the initial condition 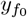 that quickly diminishes. (c) The variables *y* and *y*_*f*_ display very limited variation with respect to the initial condition *x*_0_. The population *x* exhibits the greatest sensitivity, although this also decreases over time, and the population *ϕ* again shows very little sensitivity. (d) The variables *y* and *y*_*f*_ show a notable variation with respect to the initial condition *ϕ*_0_. The variables *x* and *ϕ* exhibit a small initial variation that eventually decreases.

The results obtained for the initial conditions *y*_0_ and 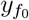 show that the variations in the different variables are very small [see Fig. 7(a,b)]. The perturbed solutions quickly stabilize, approaching the unperturbed solution. In the case of the initial condition *x*_0_, we observe some variation, especially in the variable *x*. However, all variables eventually converge to the unperturbed solution [see Fig. 7(c)]. Finally, the initial condition *ϕ*_0_ has the greatest impact on the system’s sensitivity, as observed in our results. Much larger variations occur than in the other cases, particularly for the populations *y* and *y*_*f*_ [see Fig. 7(a,b)]. This can also be interpreted as the model dynamics having a strong dependence on the initial condition *ϕ*_0_. Except for this last case, no significant variation is observed for any of the populations in the other results; all obtained values remain within orders of magnitude that support the robustness of the model.

In general, the sensitivity analyses conducted for both parameters and initial conditions allow us to conclude that our model is very stable and robust. That is, small perturbations in the fixed conditions of the model—whether initial conditions or parameter values—do not produce significant fluctuations in the solutions.

Biologically, this robustness implies that the emergence of resistant-dominated outcomes is not an artifact of fine-tuned parameter choices or experimental noise. Instead, the qualitative behavior is structurally stable across broad parameter ranges, reinforcing the relevance of the identified quasineutral manifold for real phage–bacteria systems.

## 5. Conclusions

The global proliferation of multidrug-resistant (MDR) bacteria poses a critical threat to public health by compromising the efficacy of conventional antibiotic therapies and necessitating the development of alternative antimicrobial strategies [1]. With antibiotic development pipelines lagging behind clinical needs [17], phages are regaining attention as a viable therapeutic option [40]. Clinical cases and trials have demonstrated phage efficacy against antibiotic-resistant infections [39]-[19], and engineered phages and phage cocktails are being explored to broaden host range and mitigate resistance [4]. Therapies based on combining phages with antibiotics are an important topic of current research from both the experimental [23, 33, 29, 30] and theoretical [37, 24] approaches.

Implementing phage therapy effectively requires a deep understanding of phage-bacteria dynamics, particularly to address the problem of the emergence of phage-resistant strains. Mathematical modeling provides a powerful tool to simulate and predict these interactions, supporting optimized treatment strategies [26]. *In vitro* experiments, such as those recently reported by Rodríguez et al. [36], allow estimation of key parameters like infection rates, burst sizes, and the likelihood of the emergence of phage-resistant bacteria. In the experiments carried out in Ref. [36], the initial population of *Klebsiella pneumoniae* was depleted by the phages, but after a few hours, the population of bacteria started growing again due to the emergence or overgrowth of phage-resistant strains. Previous *in vitro* experiments [7, 27, 25, 12] and ecological models [27, 3, 41] of phage-bacteria systems predict broad regimes of coexistence among phage and bacteria populations. In many instances, co-culturing of phage and bacteria together leads to the emergence of persistent coevolution, maintaining their populations in the long term through the so-called Red Queen dynamics [12, 22, 2].

In this work, we have analyzed the model introduced in Ref. [36], which was calibrated with *in vitro* data on bacteria-phage dynamics. Here we have studied the equilibria and bifurcations that govern coexistence and clearance scenarios for the phage-resistant strain(s). We have identified coexistence of phage-resistant bacteria with phages by means of a degenerate normally hyperbolic invariant manifold (NHIM). This degenerate NHIM is indeed a line of quasi-neutral equilibria with an unstable and a stable part. This NHIM, in the deterministic approach analyzed here, involves different initial conditions attracted toward different regions of the stable line, i.e., non-chaotic sensitivity to initial conditions. The presence of quasineutral manifolds has been described in other models in theoretical biology. For instance, lines or curves of equilibria have been identified in prey-predator models given by differential [14] and partial differential [16] equations, in Lotka-Volterra competition models [14, 28], in strains’ competition models of disease dynamics [20], in models of RNA genome replication [38], and in host-parasite systems [18]. Other dynamical systems have been shown to be governed by higher-dimensional degenerate NHIMs involving planes filled with equilibria. These neutral surfaces have been characterized in predator-prey models with Holling type III functional responses [15] and in socio-economical models [5] and, more recently, in a mathematical model for betacoronavirus between-cell infections [31].

Summarizing, our work provides insights into simple phage-bacteria dynamics by analyzing a model successfully calibrated with *in vitro* experimental data for *K. pneumoniae* with phages. We have described a coexistence scenario governed by non-hyperbolic equilibria, opening new avenues to understand the role of these manifolds in real systems. Despite their widespread presence in models of biological systems, these manifolds lack experimental evidence. Future research should also investigate phage-bacteria quasineutral dynamics considering stochasticity. In this sense, previous works described stochastic dynamics as relevant determinants of asymptotic behaviors with quasineutral manifolds [28, 20, 38].

Our results suggest that experimental measurements targeting trajectories along quasineutral manifolds—rather than single equilibrium states—may be crucial for understanding and controlling phage resistance. More broadly, identifying non-hyperbolic dynamical structures in data-driven biological models may reveal hidden constraints on therapeutic optimization that are invisible to purely numerical or equilibrium-based analyses.

## Acknowledgements

This research has been funded by the State Research Agency (AEI), through the Severo Ochoa and Maria de Maeztu Program for Centers and Units of Excellence in R&D (CEX2020-001084-M). We thank the CERCA Program/Generalitat de Catalunya for institutional support. This work has also been funded by project PID2020-112835RA-I00 (IC) funded by MCIN/AEI/10.13039/501100011033 to P.D.-C. Moreover, P.D.-C. was financially supported by a Ramón y Cajal contract RYC2019-028015-I funded by MCIN/AEI/10.13039/501100011033, ESF Invest in your future. F.D. has been supported by the Spanish Research project PID2023-147461NB-I00 funded by AEI. We want to thank the hospitality of the Institute for Integrative Systems Biology and the EnBiVir Lab, which hosted J.A. during the preparation of this manuscript. We also want to thank J. Tomás Lázaro and Mariona Fucho-Rius for their useful comments.

